# Controlling motion artefact levels in MR images by suspending data acquisition during periods of head motion

**DOI:** 10.1101/230490

**Authors:** Rémi Castella, Lionel Arn, Estelle Dupuis, Martina F. Callaghan, Bogdan Draganski, Antoine Lutti

## Abstract

Head movements are a major source of MRI artefacts that hamper radiological assessment and computer-based morphological and functional measures of the human brain. Prospective motion correction techniques continuously update the MRI scanner based on head position information provided by an external tracking system. While prospective motion correction significantly improves data quality, strong motion artefacts may remain with large head motions or when motion takes place at sensitive times of the acquisition. Here we present a framework that allows the suspension of data acquisition when head motion is predicted to have a strong negative impact on data quality. The predictor, calculated in real-time during the acquisition, accounts for the amplitude of the signal acquired at the time of the motion, thereby offering a re-acquisition strategy more efficient than relying on head speed alone. The suspension of data acquisition is governed by the trade-off between image degradation due to motion and prolonging the scan time. This trade-off can be tuned by the user according to the desired level of image quality and the participant‘s tolerability. We test the framework using two motion experiments and two head coils. Significant improvements in data quality are obtained with stringent threshold values for the suspension of acquisition. Substantial reductions in motion artefact levels are also achieved with minimal prolongation of scan time. However, high levels of motion artefacts occasionally remain despite stringent thresholds with the 64-channel head coil, an effect that might be attributed to head movement in the sharp sensitivity profile of this coil.

## Introduction

Head movement is known to lead to severe artefacts in MR images such as ghosting, blurring, geometric distortion and/or decreased signal-to-noise ratio (SNR). These artefacts hamper the diagnostic assessment of brain MRI scans (Andre et al., 2015) and have a strong impact on morphological (Reuter et al., 2015), diffusion-based (Yendiki et al., 2014) and functional (Todd et al., 2015) measures of the brain in imaging neuroscience studies. Quite naturally, the effects of head movement are biased towards non-compliant populations, which limits the use of MRI for the study of brain pathology, an essential cornerstone of clinical neuroscience.

The correction of motion effects on MRI data has seen much progress over recent years. Retrospective Motion Correction (RMC) techniques are applied post hoc on motion-corrupted MRI data (Pipe, 1999). An example of an RMC technique is the rigid-body realignment of successive image volumes in fMRI or diffusion-weighted series (Leemans and Jones, 2009; Worsley and Friston, 1995), a pre-processing procedure conducted routinely in imaging neuroscience studies. However, head motion that takes place within the acquisition of one image volume cannot be corrected by rigid-body realignment and the resulting image artefacts - e.g. ghosting, signal dropout, etc. remain in the MRI data. The correction of motion taking place within the acquisition of one image volume must be implemented on the subsets of raw data (*kspace lines*) that are used to reconstruct the final image. EPI based approaches used for DWI and fMRI allow the acquisition of whole-brain images in a few seconds. For anatomical MRI, the acquisition of one whole-brain image takes place over a timescale of minutes, making intra-volume motion correction particularly important there. RMC methods allow the pre-processing (e.g. filtering) of the motion trajectories recorded in parallel with data acquisition, enhancing the robustness of motion estimates before correction of the MRI data (Zahneisen et al., 2016). RMC also allows for the reconstruction of MRI images with and without motion correction, allowing the assessment of the efficiency of the motion correction scheme. Prospective Motion Correction (PMC) techniques act in real-time during image acquisition by adjusting the MRI scanner to account for patient movements (Zaitsev et al., 2006). With PMC methods, the acquired MRI data is already corrected for motion effects, avoiding the need for dedicated processing steps (Maclaren et al., 2013) and the correction of image artefacts which are difficult to remove (e.g. spin-history effects) (Yancey et al., 2011). While PMC techniques based on external motion tracking devices may be implemented with minimal change to the pulse sequence, they require specially-dedicated software libraries to integrate the motion information into the software environment of the MRI scanner. The availability of these libraries remains limited to-date.

Various methods are available to track head position during MRI scans. MR-based motion tracking using navigator echoes (White et al., 2010) is the most common as it does not require additional hardware. However these techniques require available time in the pulse sequence for the acquisition of the navigator lines without disruption of the steady-state. This might be difficult to accommodate when no dead time is available – as in the present study. Motion correction using active markers, i.e. small samples of MRI-visible material attached to the participant‘s head and wired to the scanner, has also been demonstrated (Ooi et al., 2009). However, such methods require multi-channel RF hardware and the insertion of dedicated blocks in the acquisition pulse sequence. Optical tracking system methods monitor passive markers to track head position during the acquisition (Godenschweger et al., 2016). In this study we use the PMC method introduced by Zaitsev et al. (Zaitsev et al., 2006), based on external optical measures of brain motion using the tracking system introduced by Maclaren et al (Maclaren et al., 2012). Tracking was performed by an optical camera mounted inside the MRI scanner bore. The position of the optical marker - attached to the participant‘s skull by a mouthpiece – was sent in real-time to the MRI host computer and MRI acquisition - gradient amplitudes, RF pulse frequency and phase, were adjusted accordingly during acquisition (Maclaren et al., 2012). Note that as for methods based on active markers, the markers must be strongly attached to the participant‘s head in order to obtain accurate tracking data. Maclaren et al reported a tracking accuracy of ˜0.01 mm in translation and ~0.01° in rotation (Maclaren et al., 2012). An fMRI study using this system has reported improvements in SNR by 30% to 40% and an increase in the numbers of significantly activated voxels by 70% to 330% (Todd et al., 2015). Improvements in the precision of relaxometry measures in the range of 11-24% have also been reported using this PMC technique (Callaghan et al., 2015).

Previous studies have pointed out that the lag time between tracking of the marker position and acquisition update (32ms, (Zaitsev et al., 2006)) can induce erroneous motion correction for fast motion, imposing an upper limit on the velocity of the head motion that can be accurately corrected (Maclaren et al., 2013; Zaitsev et al., 2006). To overcome this limitation, we introduce a framework that allows the interruption of data acquisition during periods of head motion detrimental to image quality. Data acquisition is resumed automatically by the MRI scanner following motion periods. A similar strategy has been introduced recently to allow reacquisition of data for excessive head motions (Aksoy et al., 2012; Bogner et al., 2014; Frost et al., 2016; Tisdall et al., 2012; Zaitsev et al., 2006) or motion artefacts (Benner et al., 2011; Porter and Heidemann, 2009). Here, we base our strategy on a predictor of the impact of head motion on image quality which accounts for the amplitude of the signal acquired at the time of the motion as well as head speed. This predictor allows suspension of data acquisition with optimal efficiency by continuing the sampling process during periods of strong motion when the amplitude of the acquired signal is low. Importantly, the presented framework allows the user to set the maximal amount of image degradation deemed acceptable in the MRI data. This determines the threshold for the triggering of the suspension of data acquisition and governs the trade-off between the quality of the MR images and the extension of the scan duration due to the suspension. This value can be adjusted on a study-specific basis depending on the compliance of the population of interest and the desired level of image quality.

## Materials and methods

### Data acquisition

Data were acquired on five participants on a 3T Siemens Prisma MRI scanner (Erlangen, Germany) using a 64 channel and a 20 channel head-neck coil and a custom-made 3D fast low angle shot (FLASH) acquisition (Lutti et al., 2014; Weiskopf et al., 2013). The kspace trajectory of the FLASH sequence was Cartesian centered (i.e. the centre of kspace was acquired half-way through the acquisition). The image resolution was 1.5mm^3^, the matrix size was 160x150x120 and the acquisition time was 7min21s per image volume. The repetition time (TR) was 24.5ms. Eight echo images were acquired following each radio-frequency (RF) excitation with echo times TE ranging from 2.34ms to 18.72ms in steps of 2.34ms.

Prior to the motion experiments (see below), subjects were requested to remain still. During this period of no motion, data were acquired with proton density (PDw), T1 (T1w) and Magnetization Transfer (MTw)-weighted contrast (corresponding RF excitation flip angles: 6°, 21°, 6° respectively) (Weiskopf et al., 2013). BÍ mapping data was acquired according to (Lutti et al., 2012, 2010) and used together with the FLASH image volumes to compute quantitative MT maps offline according to (Helms et al., 2008). PDw data only was acquired for each condition of the motion experiments. Data acquisition was repeated across multiple scanning sessions for each motion experiment, with and without suspension of data acquisition and with different head coils. The subjects were taken out of the MRI scanner between sessions and scanner calibration (e.g. shimming) was only conducted at the start of each session.

Prospective correction of head motion (Zaitsev et al., 2006) was used for all acquisitions conducted in this study. Adjustment of the MRI scanner components was implemented according to the latest head position information available from the camera prior to each RF pulse (frequency=1/TR). Note that RF excitation was maintained during periods of suspension of data acquisition in order to preserve the steady-state equilibrium of the magnetization. Data acquisition was suspended when the marker was out of the field of view of the camera to ensure optimal motion correction of all data points.

### Marker of image degradation

Maps of the MRI parameter R2* were computed from the PDw data acquired in the motion and no-motion conditions. The R2* maps were obtained by voxel-wise linear fitting of the log of the signal amplitudes across echoes, as described in (Callaghan et al., 2014; Weiskopf et al., 2013). Maps of the standard deviation of the residuals ε (standard error of the R2* estimates) were calculated as measures of the goodness of fit of the mono-exponential model. When low levels of motion artefact are present in the MRI data, R2* values are known correlates of iron and myelin concentration in brain tissue (Cohen-Adad et al., 2012; Langkammer et al., 2010; Stüber et al., 2014) and therefore of considerable interest to neuroscience studies. Head motion increases the spatial variability of R2* maps, and region-specific coefficients of variation of R2* have been shown to characterize image degradation due to head motion (Callaghan et al., 2015). Building on this idea, we used estimates of the standard deviation of the R2* values in white matter (SDR2*) as markers of motion artefact levels in the acquired data. The white matter masks were calculated by segmenting the MT maps using SPM‘s Unified Segmentation (Ashburner and Friston, 2005) and including voxels with a minimal white matter probability of 0.95. The white matter masks were eroded by 2 voxels to reduce the contribution of regions of inhomogeneous B0-field to the SDR2* estimates. This white matter mask was also used to calculate the mean of the fitting errors *ε* in the white matter.

### Motion experiment - Experiment 1

The volunteers were instructed to move during periods of 44s, i. e. 10% of the image acquisition time. The motion experiments were conducted using 5 conditions, with different onset times of the motion period (0, 44, 88, 132 and 176s) covering different regions of the 1^st^ half of the acquisition (i.e. the 1^st^ half of k-space). **Experiment 2** – The volunteers were instructed to move during periods of 3 seconds (‘*jerks*‘) to replicate motion behaviours reported from epileptic patients (Lemieux et al., 2007). Data acquisition was repeated for 4 conditions, with 10, 20, 30 and 40 jerks per acquisition. For both motion experiments, the order of the motion conditions was randomized. For experiment 2, the onset times of the jerks were also randomized: for a given condition (number of jerks), the time-course of the motion therefore varied between repetitions of the experiment across volunteers and with/without acquisition suspension.

For both experiments 1 and 2, the participants were free to adopt any motion behaviour that they preferred: no instructions were given to them on the type of motion that they should carry out (speed, amplitude, amount of translation and rotation, periods of rest…). Experiment 1 was conducted with the 64 channel head coil. Experiment 2 was performed using both the 64-channel and 20-channel head coils. Experiments 1 and 2 were first conducted with no interruption of data acquisition (but using the PMC capability) to study the relationship between the time-course of the motion and the degradation of the quality of the R2* maps (as indicated by the SDR2* values) despite conventional prospective motion correction. From this relationship, a predictor of the impact of motion on image quality was identified that could be calculated from the motion trajectory alone. Both experiments were then repeated with interruption of data acquisition when different threshold values of this predictor were exceeded. Note that one subject did not take part in the experiments with acquisition suspension.

### Predictor of motion impact

To obtain a predictor of image degradation from motion trajectories, we follow the framework introduced by Todd et al. (Todd et al., 2015). They highlighted the importance of the amplitude of the signal acquired at the time of the motion in developing aggregate measures of motion impact over the whole duration of the acquisition. The ‘encoding-weighted integrated motion’ metric *Mew* is defined as:

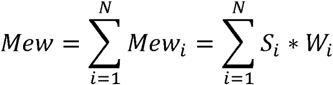

where *i* is an index over the acquired k-space lines. For the FLASH acquisition used here, the total number of acquired lines *N* is the product of the number of phase-encode steps in the phase and partition (secondary phase-encoded) directions. *W_i_* is the weight attributed to the contribution of the ith k-space line to Mew – the aggregate measure of motion impact over the acquisition of the whole 3D volume. In line with (Todd et al., 2015), we take *W_i_*, as the norm of the ith acquired line, calculated by taking the Fourier Transform of the magnitude image and summed over all readout points. Figure 1A shows an example set of weights for the acquisition used here. These weights show high/low-frequency variations due to the inner(partition)/outer(phase) phase-encoded directions, sampled using a Cartesian trajectory. *S_i_* is an index of the speed of the head at the time of acquisition of the *i*th k-space line, calculated as in (Todd et al., 2015):

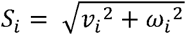

where *v_i_* and *ω_i_* are the translation and rotation speeds (in mm/s and deg/s respectively) of the centre of the field-of-view of the encoding box due to the real-time correction of head motion. Figure 1C shows the time-course of Mew, calculated from the weights and head speeds shown in figure 1A and 1B. Because the signal amplitudes, or weights, vary by several orders of magnitude throughout the acquisition (see figure 1A), a given head speed may lead to values of Mew, that may vary by several orders of magnitude too, depending on the time of occurrence of head motion. This forbids the use of a unique threshold value of Mew for the whole acquisition to detect head motion and trigger the suspension of the acquisition: a value suitable for the periphery of kspace would prevent data acquisition at the centre of kspace and the acquisition would never terminate. Conversely, a suitable value for the centre of kspace would allow for data acquisition in the periphery for any large value of the head speed. Instead this threshold value should be dependent on kspace location, considering <u>deviations</u> of Mew_i_, ΔMew_i_,) from its values when no head motion is present:

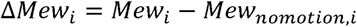

**Figure 1:**
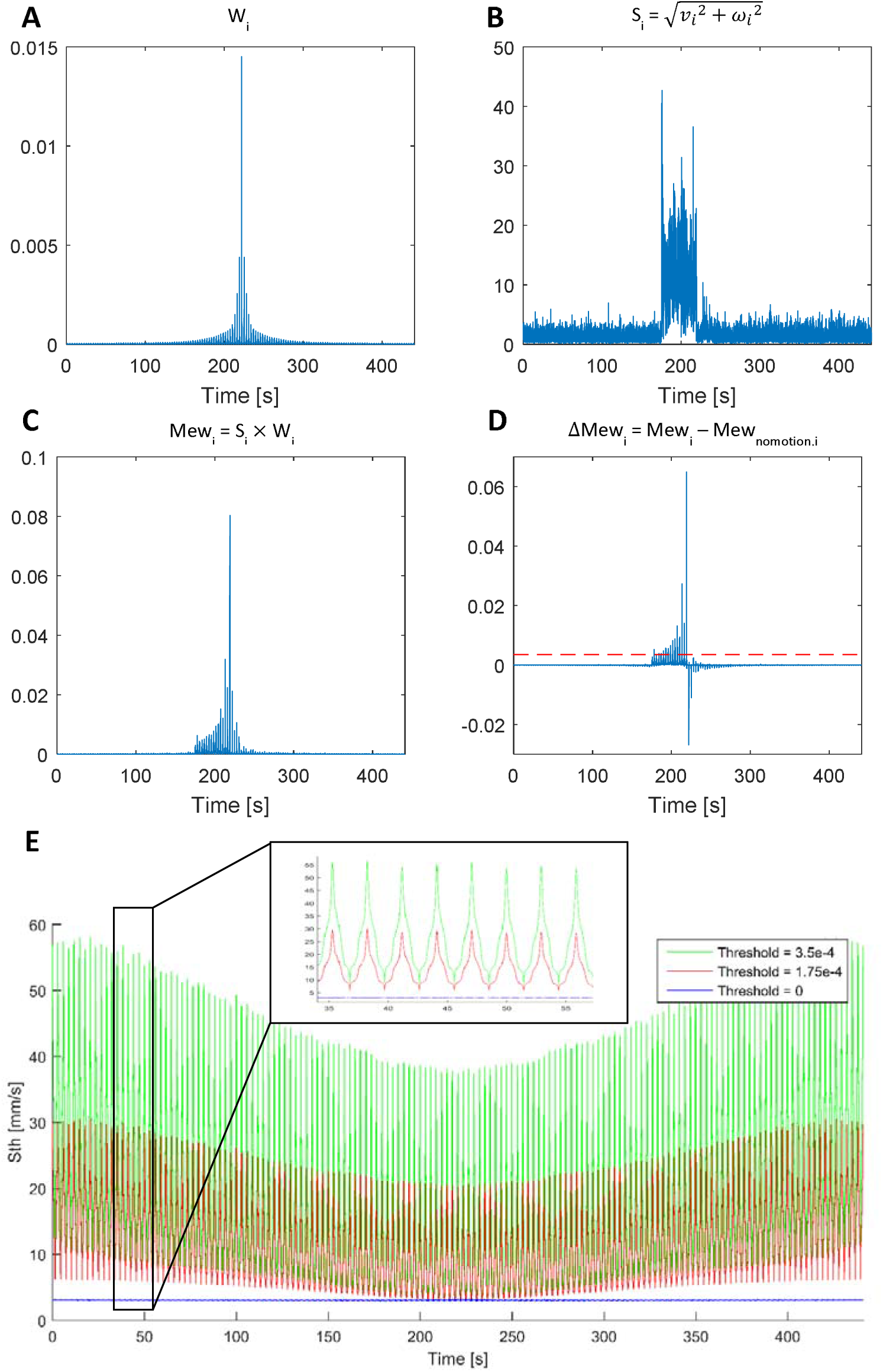
**A**) Example time evolution of the amplitude of the acquired signal (‘Weights‘) for the double phase-encoded Cartesian sampling used in the acquisitions. **B)** Example time evolution of the head speed at the centre of the Field-Of-View during an acquisition. C) Weighted motion Mew, resulting from the point-wise multiplication of A and B. **D)** Time evolution of ΔMew, the difference between Mew and reference values obtained under no motion. Values of ΔMew_i_ > 0 highlight time points where significant head motion took place. The red dashed line is an example choice of value for ΔMew_th_, the threshold ΔMew value used to suspend data acquisition. **E)** Time evolution of the threshold head speed value used for the suspension of data acquisition, equivalent to a constant ΔMew_th_ for all the acquisition (red dashed lines in D)

Here, Mew_nomotion,i_ is the value of Mew_i_, when no head motion takes place. Significant motion can then be identified in real-time for suspension of data acquisition as yielding values of ΔMeW_i_ above zero, regardless of the occurrence of the motion in kspace (see figure 1D).

The value of Mew, is virtually never zero - even with the most compliant subjects - due to noise in the estimation of the marker position/orientation and head motion of physiological origin (e.g. heartbeat and respiration). The former represents an intrinsic limit of the PMC system (in the order of a few microns, (Maclaren et al., 2012)) and the latter can be accurately corrected by the PMC system and does not need to be addressed here. Rather, it is the large increase in Mew, likely to yield significant degradation in image quality, which we seek to identify during the scans to trigger the suspension of the acquisition. Our definition of Mew_nomotion,i_ should therefore account for all possible values of S_i_, encountered in the presence of noise and for the variability in the weights across a population. Therefore, Mew_nomotion,i_ was computed from:

1. A set of reference weights obtained from PDw and T1w images acquired in 10 highly compliant participants (same acquisition sequence and image resolution as the current study) to account for the variability in weights across a population. These 10 participants (5 males, age mean/standard deviation: 37.5/13.8) were independent from those involved in the motion experiments.
2. Samples of *S* obtained from three motion time courses of five compliant participants representative of all values of S due to noise in the marker position and patient‘s physiology.

Multiple sample values of Mew_nomotion,i_ were obtained by combination of the reference weights with all possible permutations of the S samples. The reference Mew_nomotion,i_ values were obtained by taking the mean across the sample values of Mew_nomotioni,i_ and adding three standard deviations.

During image acquisition, the instantaneous value of Mew, was calculated from the product of the instantaneous head speed with the corresponding weight, calculated by taking the mean across the reference weights. Periods of motion are identified in real-time as those yielding values of ΔMeW_i_ above zero (see figure 1D). From these values, an aggregate measure of motion degradation over the entire acquisition can be calculated using:

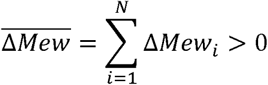

The aggregate measure of motion degradation 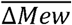 is the sum of all the values of ΔMew_i_ above zero encountered during the acquisition.

Note that suspending data acquisition based on the value of ΔMewi is equivalent to suspending the acquisition based on the head speed according to: 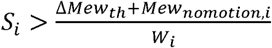 (Figure 1E). With ΔMew_th_ = 0, the suspension is most stringent: any head speed beyond that expected from physiological noise leads to suspension of data acquisition, regardless of the location in kspace. With ΔMew_th_ > 0, the threshold value of the head speed, S_th_, depends on the amplitude of the signal acquired at the time of the motion (see figure 1E): S_th_ is high/low when the amplitude of the acquired signal is low/high. Note that to avoid excessive additional scan times in case of continuous participant motion, the total duration of the suspension period was not allowed to exceed the nominal duration of the scans. Upon reaching this limit, the scanner was required to resume scanning without further suspension, leading to a doubling of the acquisition time. Note that this situation was never encountered while conducting the experiments described here.

In the 1^st^ part of this study, we show the relationship between the quality of the acquired data and the values of 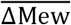. This relationship allows the use of 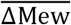 as a predictor of the impact of head motion on image quality. In the 2^nd^ part of this study, we use this predictor to monitor, in real-time during data acquisition, the impact of head motion on data quality and pause the acquisition when the predicted degradation exceeds a pre-determined threshold.

### Real-time suspension of data acquisition during periods of motion

The relationship between image degradation (SDR2*) and 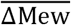 establishes the latter as a predictor of the impact of the motion time course on the quality of the data. Next, we aim to use this predictor, which is a summary metric over the whole acquisition, to trigger the suspension of the acquisition of individual data points during periods of motion. While in principle the quality of an MRI image cannot be predicted from a single time-point alone, we consider the maximum tolerable image degradation and distribute this value over the longest likely duration of the motion periods. According to motion behaviours reported from epileptic patients (Lemieux et al., 2007), a maximum of 49 jerks may be expected over the duration of the acquisition of one image volume in the present study (7min21s). This corresponds to a maximum of 147 seconds of movement during our acquisition (3s per jerk), equivalent to the acquisition of 147/TR=6000 k-space readout lines. Given the maximum aggregate value of 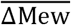 observed in our experiments (~12, see figure 2), the largest (instantaneous) value of ΔMew may be 12/6000=2e-3.Motion experiments were therefore conducted with suspension of data acquisition when ΔMew_i_ was exceeding threshold values ΔMew_th_ = 1.75e-4, 3.5e-4, 7e-4, 14e-3 and 2.8e-3, to cover all possible scenarios of patient‘s motion behaviour.

**Figure 2:**
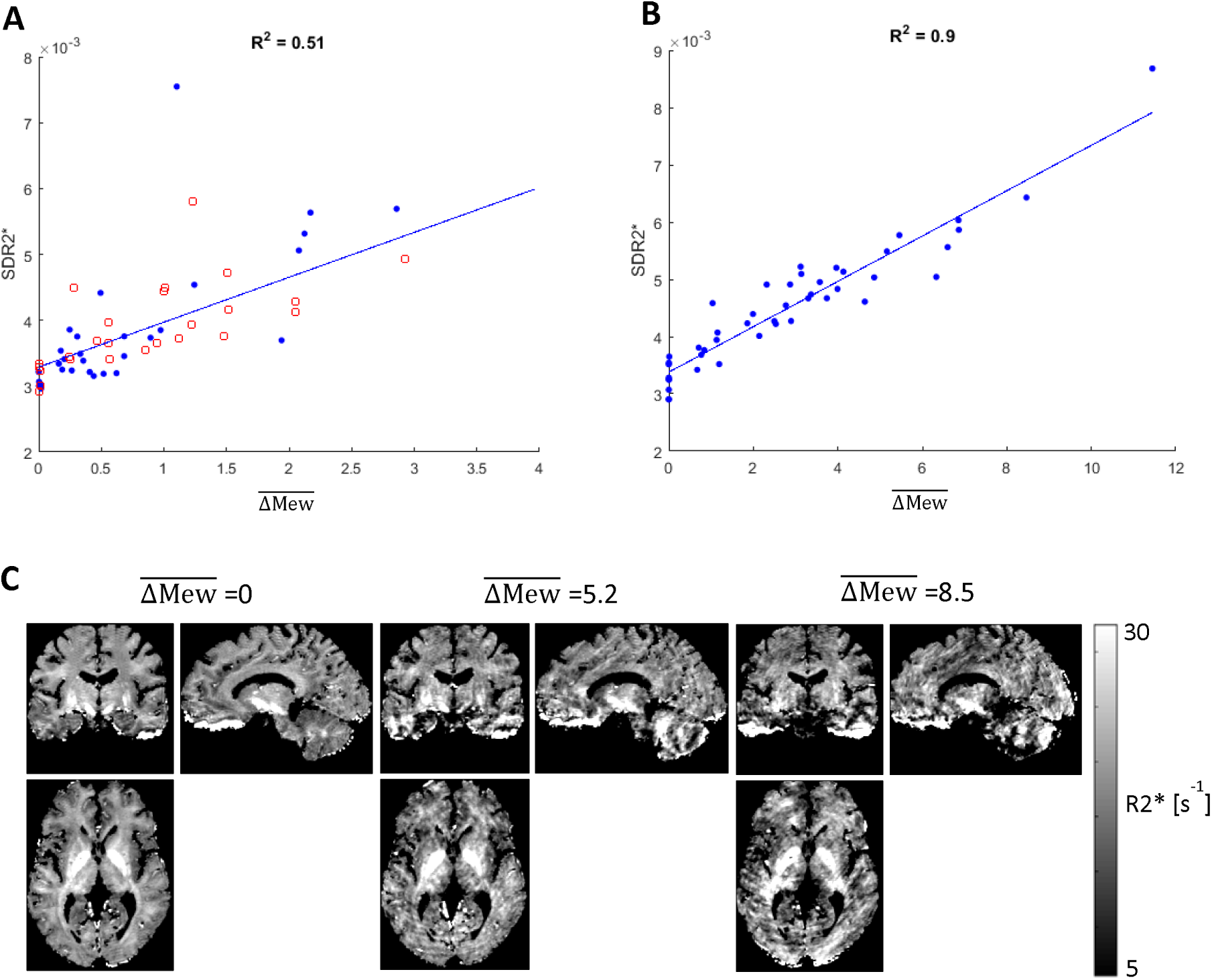
Relationship between image degradation (SDR2*) and the aggregate measure of motion 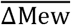 (no suspension of data acquisition). **A)** Results of experiments 1 (blue – solid circles) and 2 (red – empty circles) acquired with the 64 channel head coil. **B)** Results of experiment 2 acquired with the 20 channel head coil. **C)** Example R2* maps acquired under motion experiments leading to different 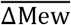 values.

### Numerical simulation

In order to further study the impact of motion on image degradation, we performed numerical simulations of head displacements in the sensitivity fields of the receive coils. We used the head translation and rotation trajectories acquired by the PMC camera during the motion experiments. These motion trajectories were retrospectively applied to the complex PDw volumes acquired during the no-motion conditions - corrected for the spatial variations in signal intensities due to the sensitivity profiles of the receive coil using the bias field correction of SPM‘s Unified Segmentation (Ashburner and Friston, 2005). Accurate replication of the image degradation observed in-vivo would have required maps of the sensitivities of the individual coil elements in the motion experiments. Because such data were not available, the separate coil-element images were acquired using a gel phantom and the same sequence as the in-vivo experiments. The spatial distribution of the phase offset of the coil elements was obtained from the voxel-wise linear regression of the phase data with the corresponding echo times. This regression included an extra regressor to account for systematic differences in phase value between odd and even echoes. Estimates of the phase and magnitude of the coil profiles outside the phantom were obtained by polynomial fitting (order 6) of the data inside the phantom.

To simulate the MRI data acquired at a given time in the acquisition, the 8 echoes of the bias-corrected PDw image were translated and rotated according to the corresponding motion data and multiplied voxel-wise by the nearest point in the receive field of the coil profiles. A 3D Fourier Transform was applied on the resulting images and the k-space line acquired at the time of the motion was extracted. This procedure was repeated for all elements of the head coils and each of the 18000 (=150*120, the total number of phase-encode steps) time frames of the acquisition. This produced simulated images of the ‘head’ moving in the receive profiles of the coil elements according to the motion trajectories obtained in-vivo. These images were combined across coil elements by a sum of square operation in image space, to match the MRI data acquired in-vivo. R2* maps were computed from the 8 simulated echo images and the SD of R2* in white matter was computed for comparison with the acquired data. Each simulation of a single MR acquisition typically lasted about two days.

These simulations were run using the motion trajectories and ‘heads’ of the 8 runs that led to the largest deviations from the linear fit in figure 4A, obtained with the 64ch head coil, and with the sensitivity profiles of the 64ch and 20ch coils. The threshold values for these 8 runs were 1.75e-4, 3.5e-4, 7e-4 and 1.4e-3. The motion information during periods of suspension was left out of the simulations.

## Results

Without suspension of data acquisition, the degradation of the R2* maps with head motion follows the same linear dependence on the parameter 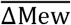 for all subjects and under both block and jerk-like motion conditions (Figure 2A, B). This linear dependence was observed for both head coils, though less robustly for the 64ch coil (R^2^ = 0.51) than for the 20ch coil (R^2^ = 0.9). Figure 2C shows examples of R2* maps for different values of 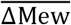. The degradation of the R2* maps increases when motion takes place near the centre of k-space (experiment 1) or with an increasing number of jerks (experiment 2). The good agreement of the relationship between 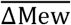 and SDR2* for both motion experiments establishes 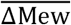 as a summary measure of motion impact, aggregated over the whole acquisition, regardless of the motion trajectories. These results, acquired with prospective correction of head motion, also constitute a baseline for the assessment of the improvements in data quality when data acquisition is suspended during periods of head motion.

Figure 3 shows example R2* maps acquired with prospective motion correction but without (A) and with (ΔMew_th_=0) (B) suspension of data acquisition, for motion conditions of experiment 1 with onset times for the block motion at: 0s, 88s, and 176s. In figure 3B, the time points corresponding to suspension of data acquisition are highlighted in red and accounted for 11%, 13% and 12% of the total number of points. The suspension of data acquisition during periods of motion leads to substantial improvements in data quality, particularly when motion takes place during the acquisition of the centre of k-space (centre of the motion trajectories). With suspension of data acquisition, the quality of the R2* maps under the motion conditions remains close to that of the R2* maps acquired without motion, even when motion takes place during the most motion-sensitive parts of the acquisition.

**Figure 3:**
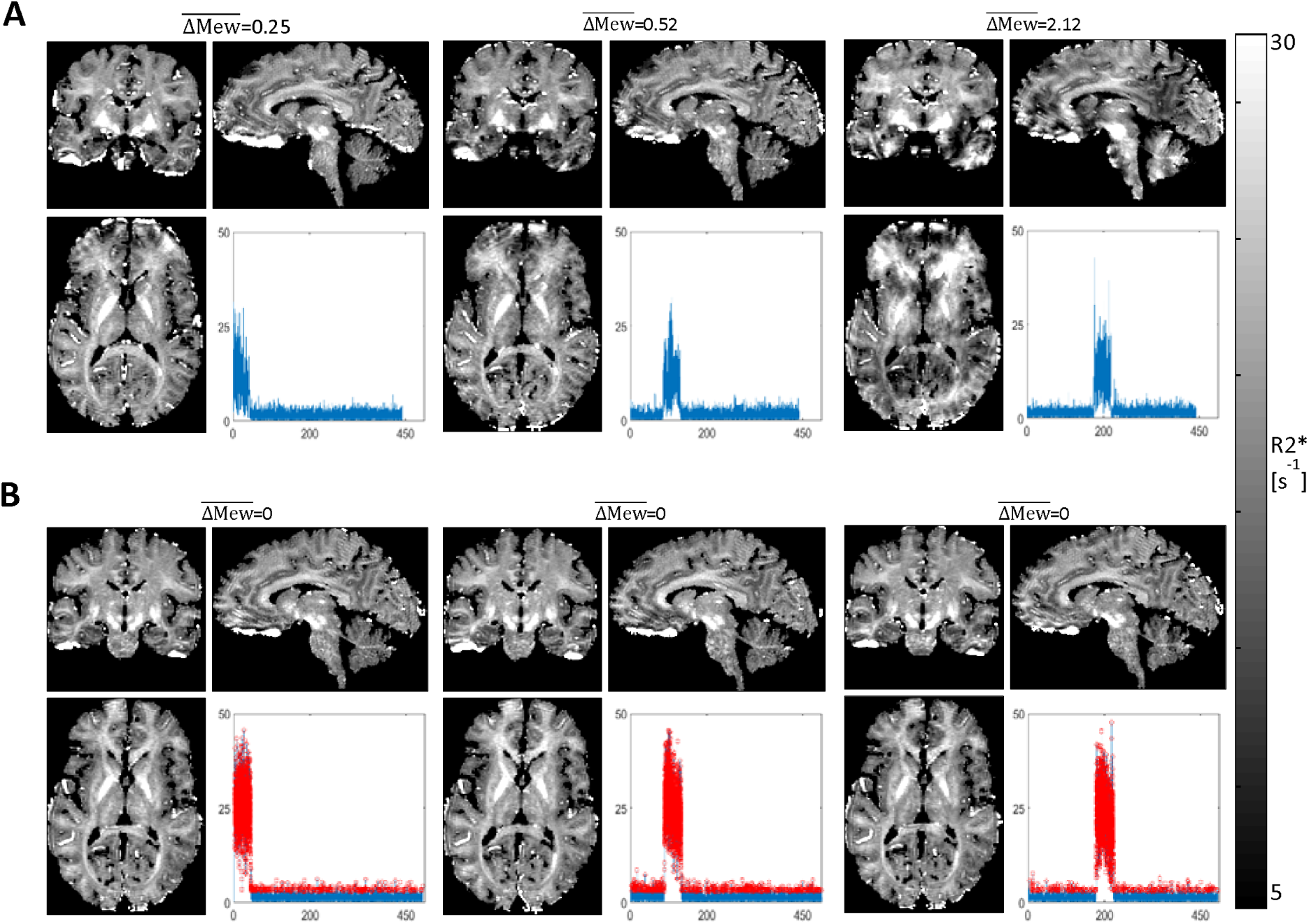
Example R2* maps acquired without (A) and with (B) suspension of data acquisition during periods of head motion. In B, the threshold value used for suspension of the acquisition was 0 and the points of suspended acquisition are highlighted in red (11%, 13% and 12% of the total number of points). The aggregate measures of motion impact – calculated from the motion trajectories – are indicated on top of each R2* map.

Table 1 shows the increase in scan time due to the suspension of data acquisition and the corresponding values of SDR2* (experiment 1). With ΔMew_th_ = 0, optimal image quality is preserved for all motion conditions at the cost of a large increase in acquisition time (45s-50s, the duration of the motion period in experiment 1). When ΔMew_th_ = 7e-4, increasing values of image degradation are observed as the motion period takes place nearer to the centre of kspace. The additional scan duration is short when motion takes place at the periphery of kspace (e.g. condition 1) because the threshold value for the head speed is high there (see figure 1E). Reciprocally, the prolongation of the scan is long, in the same range as for ΔMew_th_=0, when motion takes place near the kspace centre due to the lower head speed threshold value.

**Table 1:** Additional scan time (in percent) due to the suspension of data acquisition and corresponding SDR2* values for the motion conditions of experiment 1.

|  |  | Condition 1 | Condition 2 | Condition 3 | Condition 4 | Condition 5 |
| --- | --- | --- | --- | --- | --- | --- |
| $\Delta Mew_{th} = 0$ | Additional time [%] | 10.4 | 11.2 | 11.4 | 11.1 | 11.1 |
|  | SDR2* | 3.01e-3 | 3.09e-3 | 2.99e-3 | 3.06e-3 | 3.22e-3 |
| $\Delta Mew_{th} = 7e-4$ | Additional time [%] | 1.6 | 3.9 | 4.3 | 6.8 | 8.4 |
|  | SDR2* | 3.16e-3 | 3.27e-3 | 3.32e-3 | 3.34e-3 | 3.48e-3 |

Figures 4A and 4B show the effect of suspending data acquisition for ΔMew_th_ values of 3.5e-4, 1.4e-3 and 2.8e-3. For a given motion condition, the maximum 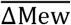 and SDR2* values tend to decrease when a lower ΔMew_th_ is used, illustrating the gain in image quality with decreasing threshold. As for the experiments without suspension of data acquisition, the robustness of the linear dependence of SDR2* on 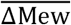 is less pronounced for the 64ch coil (R^2^ = 0.46) than for the 20ch coil (R^2^ = 0.81). Figure 4C shows the R2* maps with the highest SDR2* for each threshold value, acquired with the 20ch coil and experiment 2. Without interruption of the acquisition, the poor quality of the R2* map makes it practically unusable for neuroscience applications. The quality of the R2* maps increases as ΔMew_th_ is reduced. For ΔMew_th_=3.5e-4, the quality of the R2* maps (SDR2*=4.35e-3; 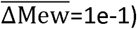 is close to that obtained under the no motion condition (SDR2*=3.08e-3; 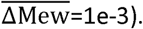

**Figure 4:**
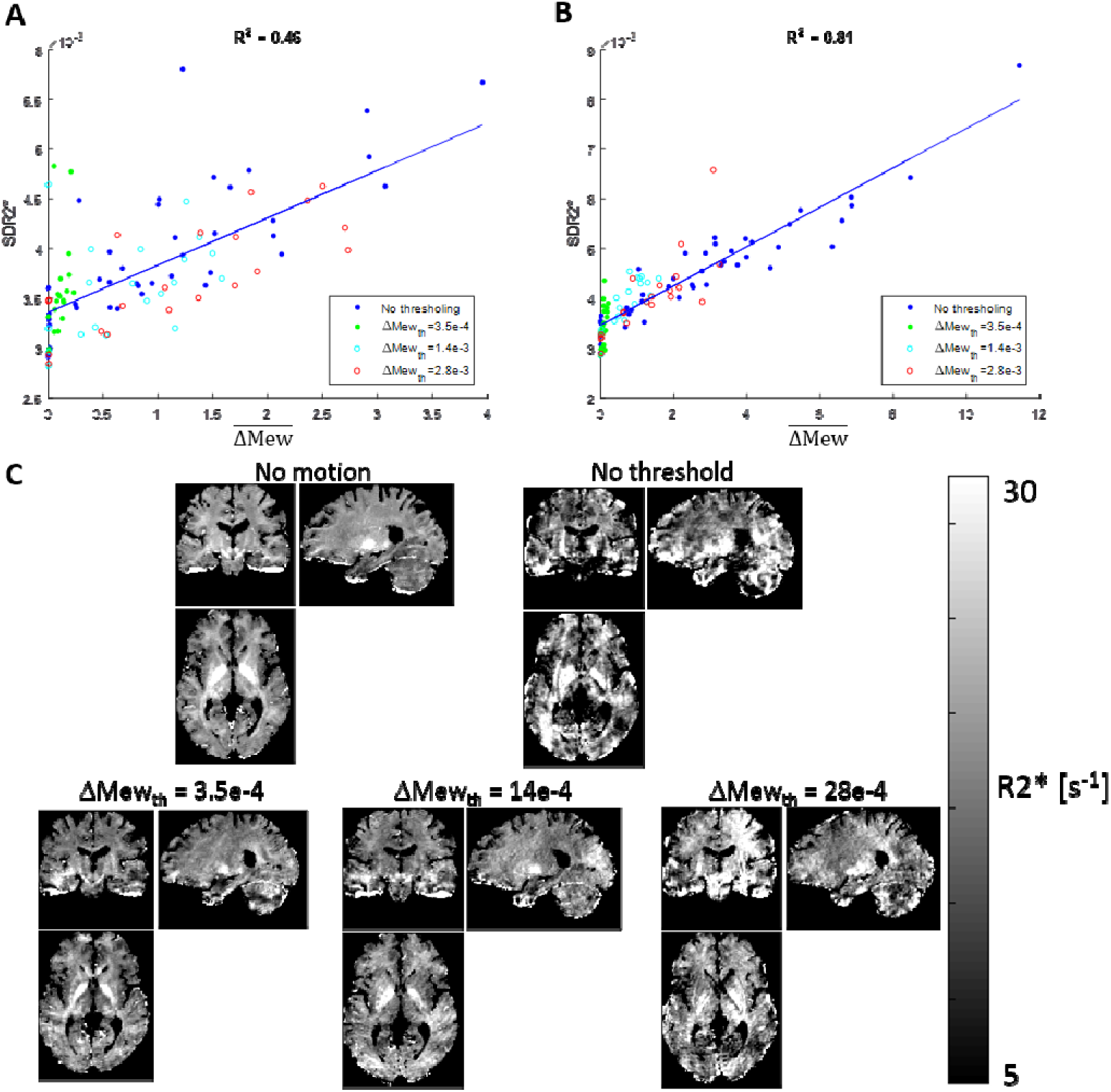
Relationship between image degradation (SDR2*) and the aggregate measure of motion, for experiment 2 with and without acquisition suspension. Data was acquired with the 64ch (**A**) and 20ch (**B**) head coils. **C**) R2* maps corresponding to the highest SDR2* values obtained for each threshold value of the acquisition suspension with the 20ch head coil.

Figure 5A and 5B show the relationship between the goodness of fit measures (residuals) and the aggregate measure of motion 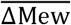 for different ΔMew_th_ values of 3.5e-4, 1.4e-3 and 2.8e-3. Without suspension of the acquisition, *ε* follows a similar linear dependence on 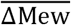 as SDR2* (figure 4). For low values of ΔMew_th_, the values of the fitting errors tend to remain in the vicinity of the values of the no-motion condition. The range of ε values increases with increasing ΔMew_th_, illustrating a decrease in the robustness of the fit, but remains smaller than those obtained without acquisition suspension. Figure 5C, shows the maps of the residuals corresponding to the highest SDR2* for each threshold and illustrates these observations.

**Figure 5:**
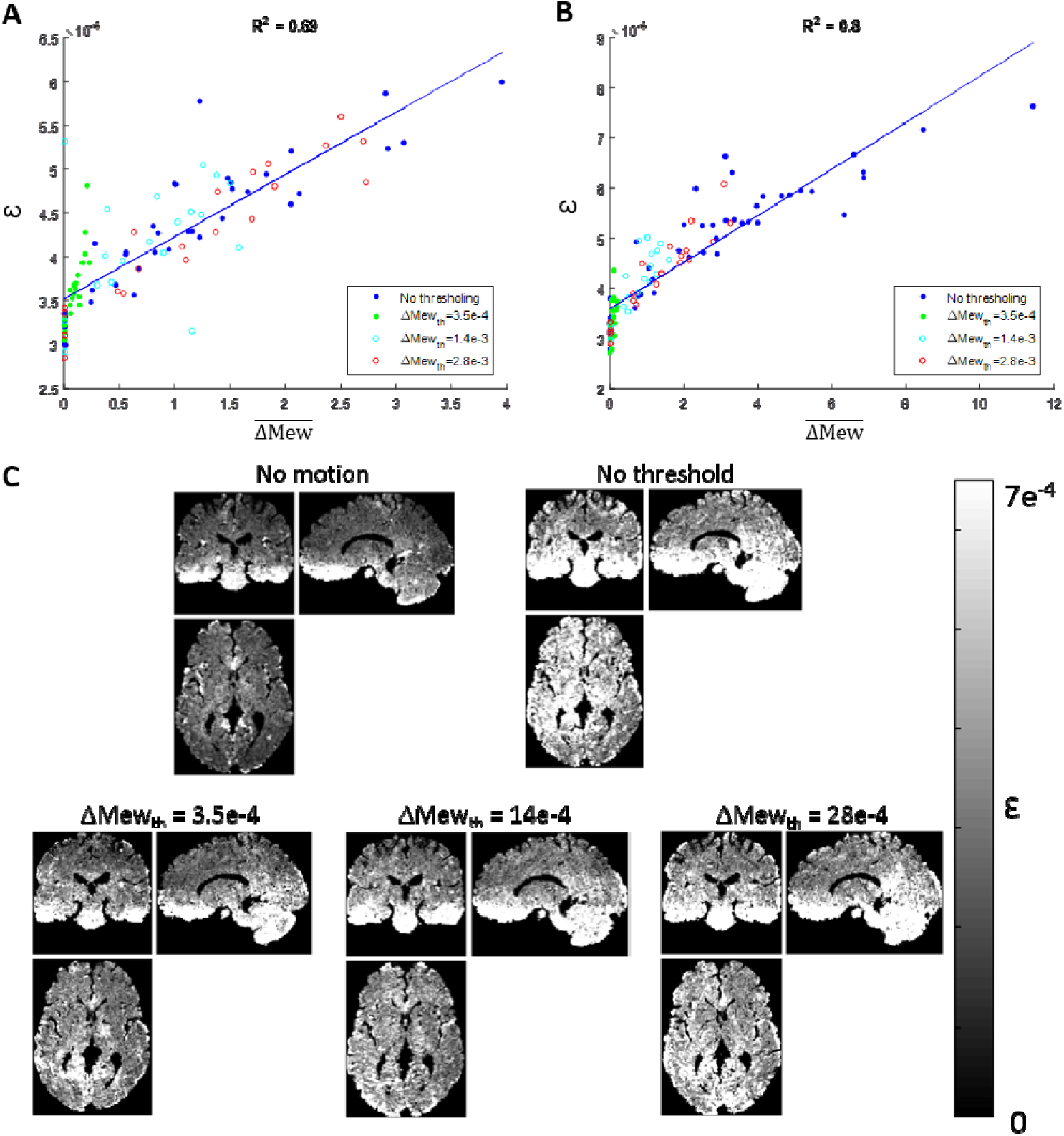
Relationship between mean residuals of the R2*fit (0) and the aggregate measure of motion, for experiment 2 with and without acquisition suspension. Data was acquired with the 64ch (**A**) and 20ch (**B**) head coils. **C**) Residuals maps corresponding to the highest SDR2* values obtained for each threshold value of the acquisition suspension with the 20ch head coil.

*Numerical simulations*

The simulated SDR2* values – indicative of image degradation due to motion - were systematically higher for the 64ch than the 20ch head coil, for the same motion trajectories (Figure 6A). For the 6 data points where the SDR2* of the acquired data was higher than that of the simulated data, the simulations with the 64ch/20ch head coil accounted for 80%/50% of the increase in SDR2* between the no-motion and motion conditions. Figure 6B shows example R2* maps acquired and simulated with sensitivity profiles obtained from the 64ch and 20ch head coils. Although the pattern of artefacts observed experimentally could not be reproduced with the simulations, the increased level of image artefacts with the 64ch coil, as captured by SDR2*, can be observed. This suggests that the sharper sensitivity profiles of the 64ch coil are a dominant contribution to the reduced correlation between 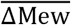 and SDR2* visible in figures 2 and 4.

**Figure 6:**
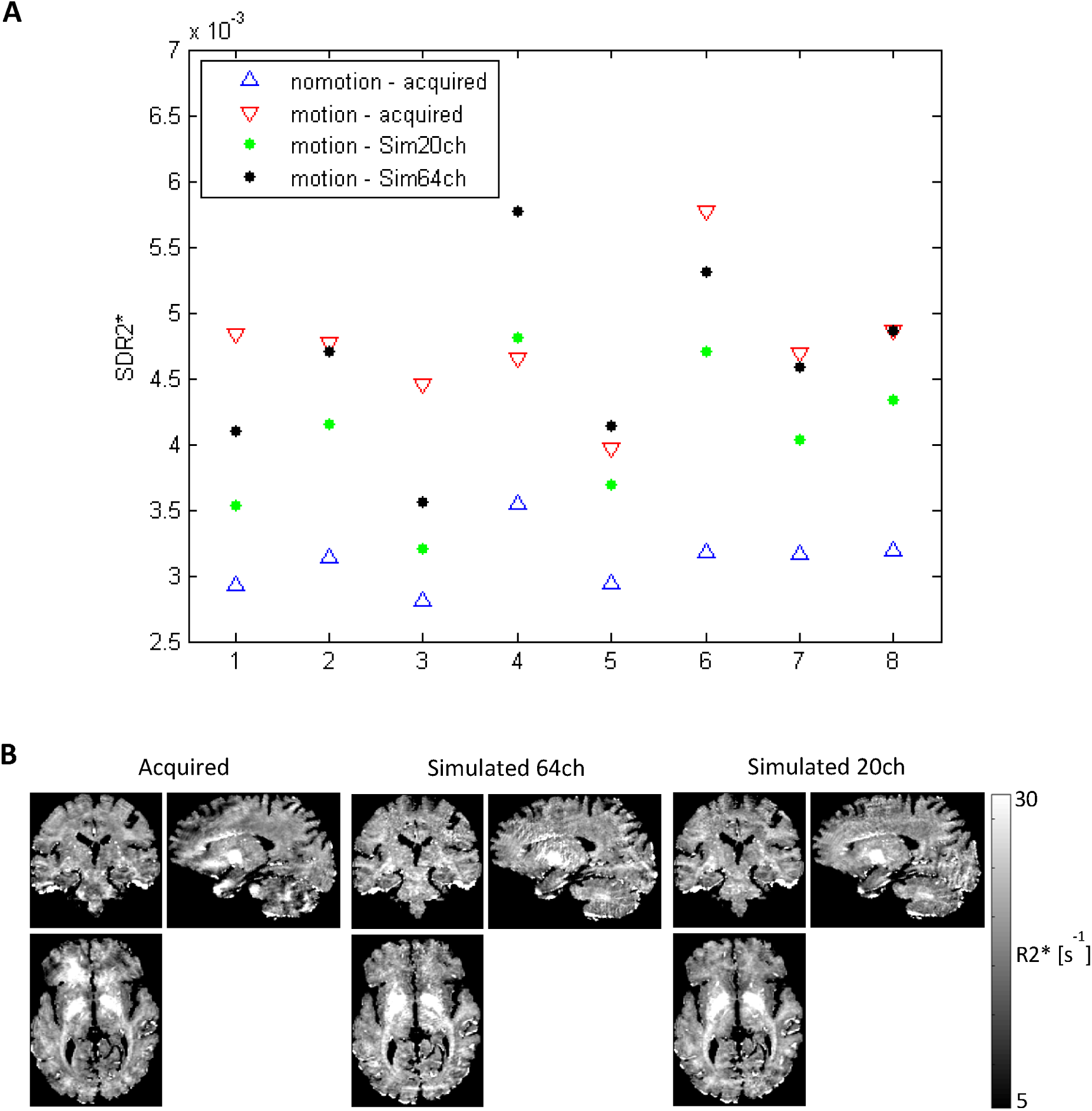
Simulation of head motion in the spatially varying sensitivity profiles of the 64ch and 20ch head coils. The simulation was run on the 8 motion time courses that led to large deviations from the linear behaviour shown in figure 4A. **A**) Experimental and simulated SDR2* values. The corresponding SDR2* obtained experimentally under the no-motion condition are shown for reference. **B**) Example of acquired R2* map and corresponding simulated R2* maps with the 64ch and 20ch profiles.

## Discussion

In this study, we present a fraMework to improve the quality of MRI data when large head motions take place that cannot be accurately compensated by PMC correction. The fraMework relies on a prospective system for motion correction based on an optical camera that tracks head position during the scans and provides this information to the MRI scanner for adjustment in real-time (Maclaren et al., 2012). While the high level of performance of this system has been demonstrated (Callaghan et al., 2015; Todd et al., 2015), large motion-related artefacts remain in the acquired data with motion behaviours representative of patient populations. The proposed fraMework addresses this issue by suspending data acquisition during periods of motion detrimental to data quality. We present an aggregate measure of the motion time course during the acquisition that exhibits a high correlation with the degradation of the resulting image (figure 2). From this predictor of motion impact, a threshold parameter can be defined to suspend image acquisition during periods of motion that are likely to be detrimental to image quality (figure 3). This threshold guarantees a minimum level of data quality in the final images, given the motion characteristics of patient populations (figure 4). This threshold also governs the trade-off between image quality and the prolongation of scan time due to the suspension of the acquisition. The value of this threshold can be set by the user depending on the desired level of image quality and the patients’ tolerance for prolonged scan durations. For low threshold values, the image quality remains largely comparable to that acquired without motion, at the expense of a large increase in scan time when long periods of motion take place. Note that to ensure completion of the scans in cases of continuous participant motion, the maximum total duration of the suspension was set to the nominal duration of the scans. Upon reaching this limit, the acquisition was designed to resume scanning without any further suspension of the acquisition, leading to a doubling of the scan time. While this feature was implemented as a safeguard, this situation was never encountered in practice. Beside head speed, the predictor of motion impact accounts for the time of occurrence of the motion in k-space. As a result, periods of strong motion may be tolerated at innocuous times of the image encoding, in order to minimise the increase in acquisition time at a minimal cost in terms of image quality (table 1)·

The relationship between the aggregate measure of motion and image degradation was examined for two types of head movements: continuous motion periods of 44s in duration with pre-defined onset times (experiment 1) and short motion periods of 3s in duration (‘jerks‘) at random times during the scans (experiment 2). The comparability of the participant‘s motion behaviour between motion conditions was not a pre-requisite for this study. To the contrary, no instructions were given to the participants regarding their motion behaviours and in experiment 2, the onset times of the jerks were randomized across repetitions, enforcing a high degree of variability. Despite this variability between participants, motion experiments and conditions, the relationship between image degradation and the motion impact was found to be remarkably consistent, suggesting that the predictor of motion impact – aggregated over the whole acquisition – is robust against the variability of motion time-courses that might be encountered practically with non-compliant populations. In preliminary experiments, this correlation was found to be consistent when the TR duration of the acquisition was extended to 50ms - double the TR value used here (data not shown). This indicates that the presented relationship may hold for TR values typically used with the FLASH acquisition in practice.

For a given condition/number of jerks, the inherent variability between repetitions of experiment 2, due to the randomness of the onset times of the jerks, makes it unsuitable to estimate representative figures of the prolongation of scan time due to the suspension of data acquisition. Triggering of data suspension is indeed determined by the occurrence of the jerks in kspace. Instead, this was achieved with experiment 1 (motion periods with pre-defined onset times), for which the motion behaviour may be more safely assumed reproducible across multiple repetitions of the same condition. The results are presented in table 1. Suspension of data acquisition based on head speed only (ΔM_ewth_=0) led to systematic prolongations of scan time by the duration of the motion periods. With ΔM_ewth_=7e-4, the prolongation of scan time was seen to vary by a factor 5 depending on the time of occurrence of the motion.

The presented data shows a clear decrease in the correlation between image quality and the motion predictor with the 64ch head coil (figure 2A). With that coil, MRI data of low quality was occasionally acquired, despite low threshold values for the suspension of data acquisition (figure 4A). Upon observation of these results, we conducted numerical simulations to investigate the effect of the different sensitivity profiles of both head coils. In these simulations, artefact-free head images were moved in the spatially-varying sensitivity profiles of the head coils according to the same motion trajectories, recorded in the in-vivo motion experiments. 80%/50% of the degradation in image quality observed experimentally – as captured by SDR2* - were replicated with the simulations using the sensitivity profiles of the 64ch/20ch head coils respectively. The larger image degradations obtained with the 64ch coil, for the same motion trajectories, suggest that changes in the position of the head within the coil sensitivity profiles due to the motion is a significant source of image degradation, though parallel imaging was not used in our experiments. This is consistent with the reduced correlation between image quality and the motion predictor for the 64ch head coil (figure 2A). This effect has recently been highlighted as a significant contributor to image degradation (Bammer et al., 2007). Correction of the effect requires knowledge of the sensitivity profiles of the individual coil elements and real-time implementation of the correction during the scans on each individual acquired kspace line – which would present an insurmountable challenge. A more feasible alternative would be to extend the suspension of data acquisition to include periods where the head position is too far from its original position. Note that the mere purpose of these simulations was to investigate, upon observation of our results, the potential role of the sensitivity profiles on the reduced correlation between SDR2* and 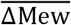 in figure 4A. The accurate reproduction of the levels and patterns of image degradation observed experimentally would have required the estimation of the sensitivity profiles of the coil elements in parallel with the motion experiments. Instead, the coil element images used in our simulations were obtained from phantom data. Also, only the changes in head position relative to the coil profiles were accounted for by the simulation and other contributions to image degradation (e.g. loading of the RF coil, orientation dependence of the MRI signal (Yarach et al., 2016), gradient non-linearity (Yarach et al., 2015), transverse coherences,…) were not accounted for. However simplistic, the numerical simulations indicate stronger image degradation for the 64ch head coil, suggesting an effect of the coil sensitivity profiles consistent with the results presented in figures 2 and 4 and in-line with recent literature (Bammer et al., 2007).

While the presented fraMework for suspension of data acquisition during periods of detrimental head motion was designed for FLASH, it might be extended to other types of image acquisitions. However, suspension of data acquisition might in general not be recommended for functional MRI (fMRI), where the synchronicity of image acquisition and functional stimulation is paramount. The key in implementing the proposed fraMework for different acquisitions lies in associating incoming head positions with specific locations in kspace. In the context of EPI acquisitions, this would require new motion information at the frequency of the EPI readout sampling (~200Hz) – out of range of our system – and would also require careful implementation to prevent the occurrence of image ghosting. However an implementation based on head speed only, prior to RF excitation, is likely to bring significant improvements in data quality (e.g. diffusion MRI) in non-compliant populations. The proposed fraMework is particularly well suited for acquisitions (2D or 3D) where each head motion information used for prospective correction can be associated with acquired data of similar amplitude. However the details of the implementations might require specific adjustments for each type of acquisition (e.g. FLAIR, MPRAGE,….) due to differences in contrast manipulation (e.g. inversion pulse). In particular, the detection of detrimental motion periods involved the calculation in real-time of the deviation between the instantaneous values of Mew with reference values in compliant populations (*Mew_nomotion*). In the current study, the reference datasets were taken from an adult population of FLASH datasets with PDw and T1w contrasts, consistent with the motion experiments presented here. In general, matching the reference datasets with the population of interest (e.g. children with smaller heads) and the type of acquisition used may be preferable. However, if different reference datasets lead to consistent values of Mew during motion periods, they can be safely combined using the fraMework presented here to obtain an implementation generalizable to multiple populations and acquisition types.

This study demonstrates a linear relationship between the proposed aggregate measure of motion impact and image degradation (figure 2) and illustrates the improvements in data quality when low spatial frequencies are preferentially targeted to trigger the suspension of data acquisition (figure 4). These results open the way for follow-up studies comparing the real-time data suspension presented here with other strategies to minimize motion impact. On the model of previous works aimed at mitigating physiological effects on MRI data (Bailes et al., 1985; Cho et al., 1990; Frost et al., 2014), one candidate approach might consist in adjusting the kspace trajectory of the acquisition in realtime depending on the motion behaviour. With this approach, regions of kspace with low signal amplitude (high kspace values) would be acquired during periods of head motion, with little effect on image quality. This approach would alleviate the need for suspension of data acquisition, keeping the acquisition time minimal for all motion behaviours. Another alternative, using the theoretical fraMework presented here, would be to complete the acquisition of the whole image without suspension, and to re-acquire post-hoc the k-space lines most affected by motion until a target 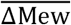 is reached. This strategy has been implemented in previous studies by identifying k-space lines corrupted by motion based on motion score (Frost et al., 2016; Tisdall et al., 2012) or magnitude/phase data (Benner et al., 2011; Porter and Heidemann, 2009). This approach would circumvent the need for assumptions on the expected amount of head motion to determine threshold values from aggregate measures of motion impact (see Real-time suspension of data acquisition during periods of motion section above), and would adjust naturally to the different motion behaviours encountered. Note that this strategy implies that the acquisition of neighbouring data points might be separated by long time intervals, which could degrade the phase consistency of neighbouring kspace lines and exacerbate image degradation (e.g. ghosting) when dynamic effects are present during the acquisition (Bailes et al., 1985; Cho et al., 1990). In light of the results presented here, changes in head position relative to the sensitivity profiles of the coil might be an example for such effects. However whether this effect is stronger with this alternative is likely to depend on the type of motion present and remains to be determined. Low-frequency scanner drift (Smith et al., 1999) might also contribute and can be minimized by conducting shimming adjustments in real-time during the scans (Bogner et al., 2014; El-Sharkawy et al., 2006; Hess et al., 2011; Keating and Ernst, 2012; Saleh et al., 2016). The identification of the optimal strategy for the mitigation of motion artefacts requires a full comparison study of these alternative strategies. The relationship between the motion impact parameter ΔMew and the image quality parameter SDR2* was shown to hold for a variety of motion behaviours and might alleviate the need for identical motion behaviours for an unbiased comparison between these strategies.

Here we use the standard deviation of R2s maps in white matter (SDR2*) as a marker of image quality. We chose this parameter because it can be computed with ease, because of its sensitivity to motion-related image degradation (Callaghan et al., 2015), and because quantitative maps of MRI parameters do not suffer from the sources of image bias that commonly affect anatomical MRI data (Weiskopf et al., 2013). Other markers of motion artefacts might be equally suitable (Pannetier et al., 2016). R2* maps are known to show a degree of variability due to the orientation of the head relative to the main magnetic field (Cohen-Adad, 2014). This orientation dependence – and its interaction with shimming - is likely to have had an impact on the estimates of SDR2* reported infigures 2 and 4, which would have reduced the correlation between ΔMew and SDR2*. The corresponding variability in SDR2*, estimated from the data acquired across multiple scanning sessions in the absence of head motion, was found to be up to 3e-4, one order of magnitude smaller than changes in SDR2* due to head motion. The results of figure 2 and 4 also include interaction effects between head position and shimming but display a remarkable level of correlation between ΔMew and SDR2* nonetheless. Note that the effect of this interaction was reduced in our analysis by eroding the white matter masks by 2 voxels prior to the calculation of the SDR2* estimates toexclude voxels near the air-tissue interface. On top of motivating the use of ΔMew for suspension of data acquisition in real-time, the linear increase in SDR2* with ΔMew also suggests that SDR2* might be used as an objective measure of data quality, potentially more robust than visual assessment. However it should be emphasized that SDR2* depends on the level of noise in the acquired data (image resolution) and on the robustness of the R2* fitting (number of echoes). It should also be noted that as R2* is a biomarker of the microstructural properties of brain tissue (Cohen-Adad et al., 2012; Langkammer et al., 2010; Stüber et al., 2014), SDR2* values might be affected by microscopic changes in the healthy and diseased brain.

## Conclusion

Here, we present a fraMework that allows the suspension of MRI data acquisition during periods of head motion. In this fraMework, head motion information is provided by a prospective motion correction system during the scans and is processed in real-time to compute a predictor of the impact of motion on the MR images. Data acquisition is suspended when excessive motion degradation is predicted. The predictor of motion impact accounts for both head speed and the amplitude of the k-space signal acquired at the time of the motion, which allows for a more efficient reacquisition strategy than one solely based on head speed. The fraMework was tested for two different kinds of motion behaviour. Suspending data acquisition during impactful periods of head motion led to significant improvements in the quality of the MRI images. The trade-off between image degradation due to motion and prolongation of scan time due to the suspension of data acquisition may be tuned depending on the desired MRI image quality and subject tolerability.

## Acknowledgments

This work was carried out on the MRI platform of the Département des Neurosciences Cliniques - Centre Hospitalier Universitaire Vaudois, which is generously supported by the Roger De Spoelberch and Partridge Foundations. AL is supported by the Roger De Spoelberch Foundation. BD is supported by the Swiss National Science Foundation (NCCR Synapsy, project grant Nr 32003B_159780), Foundation Parkinson Switzerland and Foundation Synapsis. The research leading to these results has received funding from the European Union‘s Horizon 2020 research and innovation program under grant agreement No. 720270 (HBP SGA1). The authors would like to thank Dr. David Carmichael (Institute of Child Health, University College London, UK) for helpful discussions on the movement characteristics of epileptic patients.

